## Supplementary information for "Auditory contrast gain control predicts perceptual performance and is not dependent on cortical activity"

Supplementary Figures 1-7

Supplementary Methods

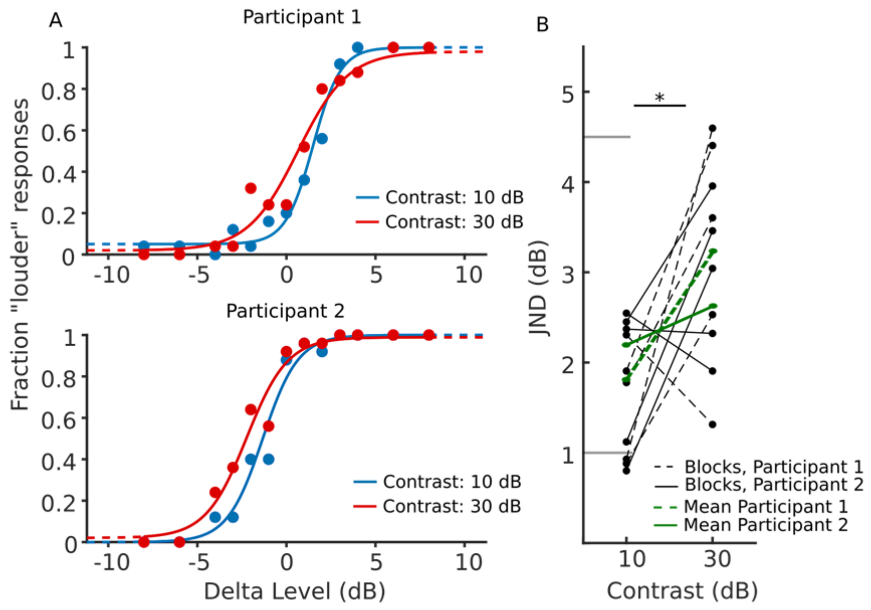


**Supplementary Figure 1. Small sound level differences between high and low contrast stimuli do not explain improved loudness discrimination performance in low contrast conditions in human listeners.** (A) Psychometric functions of two example participants in high and low contrast conditions when the overall level of the stimuli was matched. (B) Summary panel showing changes in loudness discrimination ability (as measured by the just noticeable difference, JND) between low (10 dB) and high (30 dB) contrast stimuli across blocks (black) and participants (green). The difference in JND between low and high contrast stimuli was significant (repeated measures ANOVA, treating blocks and subjects as separate random effects with contrast being the repeated measure, $p$ = 0.02). The JND increased by 49.2% (compared to 38.8% in main experiment), on average, when the contrast was increased three times (10 dB vs 30 dB).

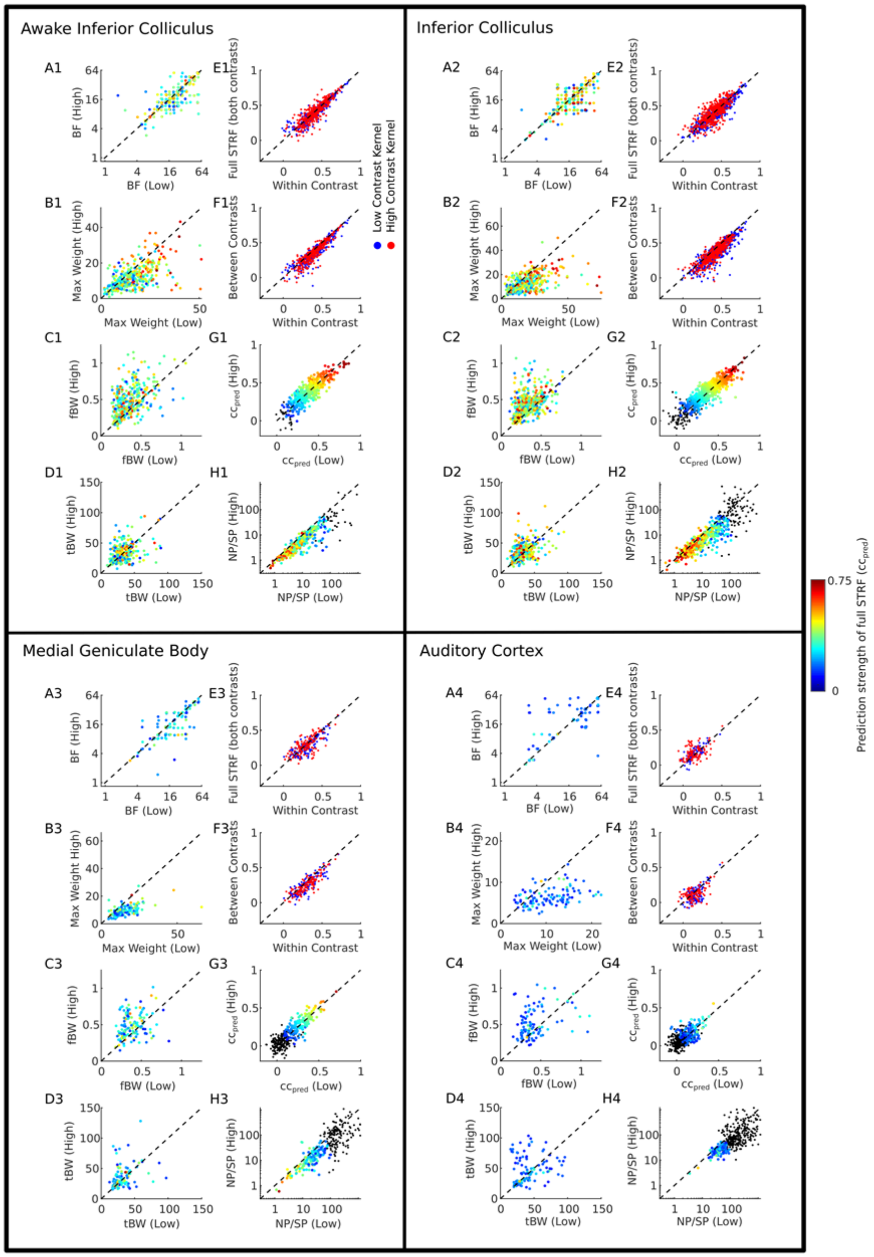


**Supplementary Figure 2. STRF changes, response reliability, and model performance within, across, and between contrast conditions in CNIC, MGBv and A1.** (A) Comparison of unit best frequency (BF), the largest value of the spectral kernel of the STRF, between low and high contrast stimuli. (B) Comparison of the value of the largest (max) weight in the spectral kernel of the STRF between low and high contrast stimuli. (C) Comparison of the frequency bandwidth (fBW), the full-width half-maximum (measured in octaves), around the BF between low and high contrast stimuli. (D) Comparison of the temporal bandwidth (tBW), the full-width half-maximum (measured in ms), around the BF between low and high contrast stimuli. (E) Prediction performance ($cc_{pred}$) of within-contrast STRF models compared to the full STRF (estimated from data from both contrasts), for high-contrast (red) and low-contrast (blue) data. (F) Prediction performance ($cc_{pred}$) of within-contrast STRF models (estimated from data of the same contrast condition), compared to between-contrast STRF models (estimated from data from the other contrast condition) for high-contrast (red) and low-contrast (blue) data. (G) Prediction performance ($cc_{pred}$) of within-contrast STRF models for low-contrast data, compared to within-contrast STRF models for high-contrast data. (H) Comparison of the ratio between noise and signal power (NP/SP) of each unit in low and high contrast conditions (see Supplementary Methods). (A-D, G-H) Color of points denotes the prediction strength (correlation coefficient) of the model on a cross-validated dataset. Black dots indicate excluded units from modelling results. $n_{CNIC}$ = 499, 13 mice, $n_{MGBv}$ = 136, 8 mice, $n_{A1}$ = 106, 10 mice, $n_{CNIC\_Awake}$ = 380, 6 mice.

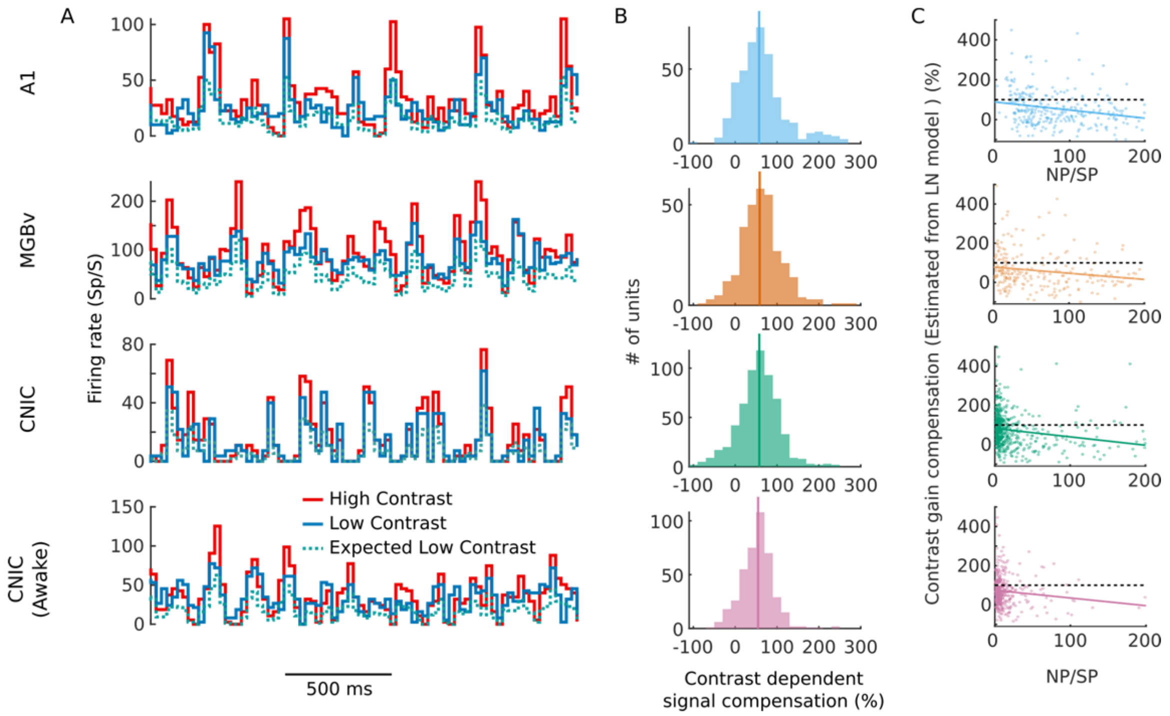


**Supplementary Figure 3. Compensated response signal (signal standard deviation) following contrast adaptation in the midbrain, thalamus and cortex, and dependence of contrast gain control (estimated using LN model) on NP/SP.** (A) Two-second PSTH snippets from example units in A1, MGBv and CNIC in anesthetized and awake mice) during high (40 dB, red) and low contrast stimulation (20 dB, blue), together with expected uncompensated low-contrast PSTHs (i.e. high-contrast PSTHs / 2). (B) Summary of signal compensation between high and low contrast (see Supplementary Methods). Units recorded in each brain region compensated for contrast, $\text{Median}_{A1}$ = 57.2% compensation, $p < 0.001$; $\text{Median}_{MGB}$ = 58.5% compensation, $p < 0.001$; $\text{Median}_{CNIC}$ = 57.6% compensation, $p < 0.001$; $\text{Median}_{\text{Awake CNIC}}$ = 54.7% compensation, $p < 0.001$; Wilcoxon sign rank tests. No significant differences were found between areas (Kruskal-Wallis test, $p = 0.30$) or between CNIC data from anesthetized and awake mice (Wilcoxon rank sum test, $p = 0.13$; $n_{A1} = 414$, $n_{MGB} = 317$, $n_{CNIC} = 603$, $n_{CNIC\_awake} = 411$). (C) Contrast gain control (as measured with LN model) as a function of NP/SP (up to NP/SP of 200 and contrast gain compensation < 500%). Solid line indicates least squares fit. In each brain region, contrast gain compensation decreased with increasing NP/SP, and the regression line intersected with the y-axis at values indicating strong contrast gain control.

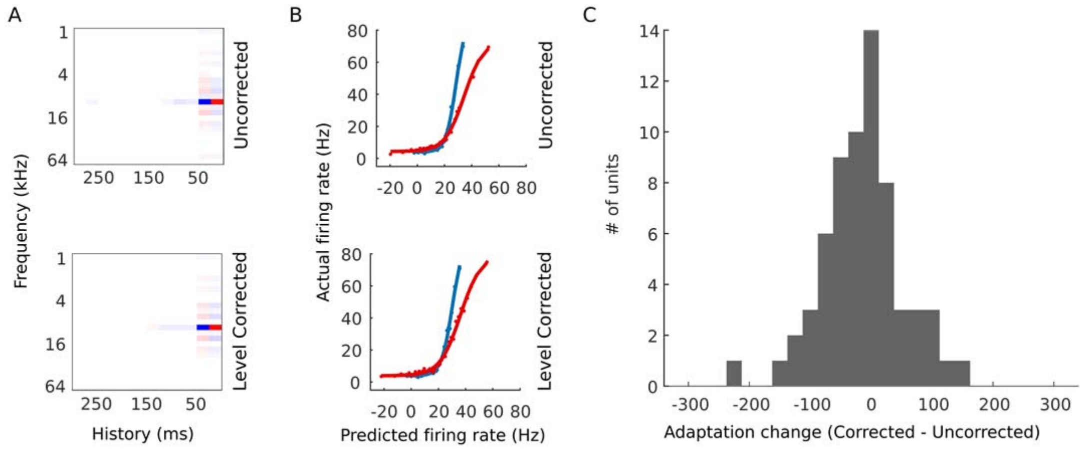


**Supplementary Figure 4. Small sound level differences between high and low contrast stimuli do not explain contrast gain control in CNIC units.** (A) STRFs from an example CNIC unit estimated using standard DRCs ("uncorrected") or DRCs whose level was reduced to account for small increase in sound level at high contrast ("level corrected"). (B) Contrast dependent output nonlinearities for an example unit, estimated from uncorrected and level-corrected DRCs. (C) Summary of difference in contrast gain control between uncorrected and level-corrected DRC stimulation. No significant difference (Wilcoxon signed-rank test, $p > 0.05$) was found.

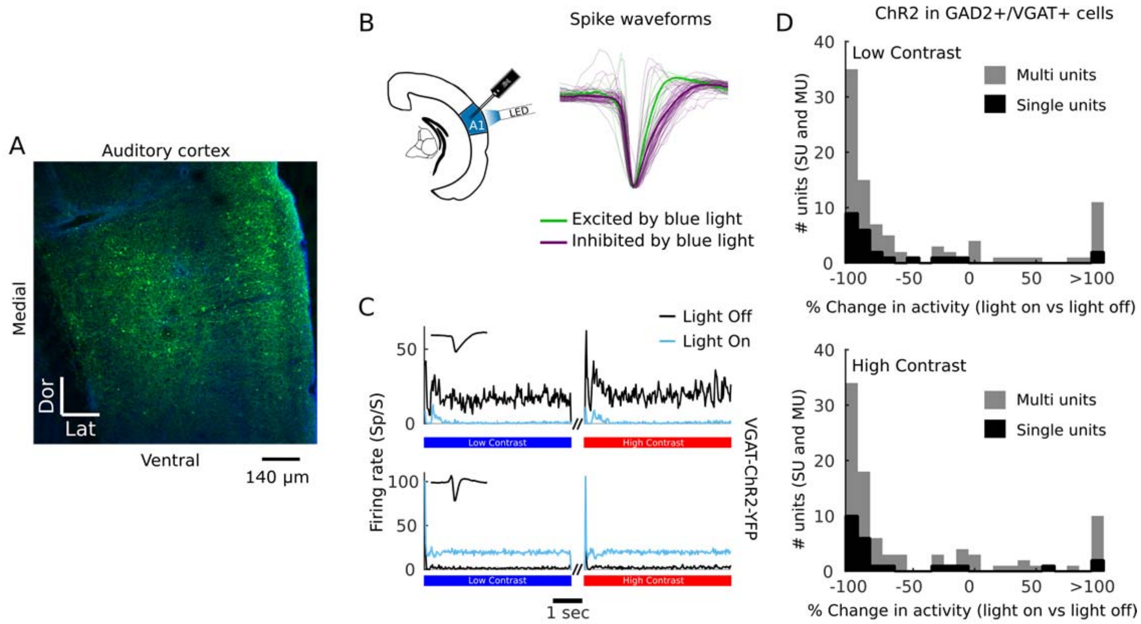


**Supplementary Figure 5. Auditory cortical silencing by driving inhibitory neurons (GAD2+/VGAT+).** (A) Confocal image of ChR2-YFP expression in auditory cortex 4 weeks after cortical injection of AAV5-DIO-ChR2-YFP in a GAD2-cre mouse. (B) Spike waveforms for units significantly excited by light stimulation (green) and units significantly inhibited by light stimulation (purple). Thin lines are individual units. Thick lines are means of individual units from each group. (C) Examples of mean PSTHs (across all trials and all high (red) or low contrast (blue) DRCs, with either light on (light blue) or off (black)) from a multi-unit inhibited by light stimulation (top) and a multi-unit driven by light stimulation (bottom). Light-dependent changes in activity were maintained for the duration of the DRC stimulation (i.e. 5 seconds). (D) Effect of light stimulation on average firing rate of single- and multi-units during low (top) or high (bottom) contrast DRC stimulation ($n = 92$, 4 mice; 1 VGAT-ChR2-YFP, 3 GAD2-cre + Floxed ChR2 virus). The activity of 68.5% of units was significantly (t-test, $p < 0.05$) suppressed at high and low contrast for the duration of the DRC stimuli by either a single 5-second light pulse (2 mice) or 5-second 40 Hz sinusoidal stimulation (2 mice). The inhibited units were mostly putative excitatory units, with regular (broad) spike waveforms. We also found that light significantly (t-test, $p < 0.05$) facilitated activity in a small subset (8.7 % of units) at both high and low contrast; these were mostly putative inhibitory units, with narrow spike waveforms. After removing units which were significantly driven by light (i.e. VGAT+/GAD2 cells), the median change in activity in auditory cortex was –88%.

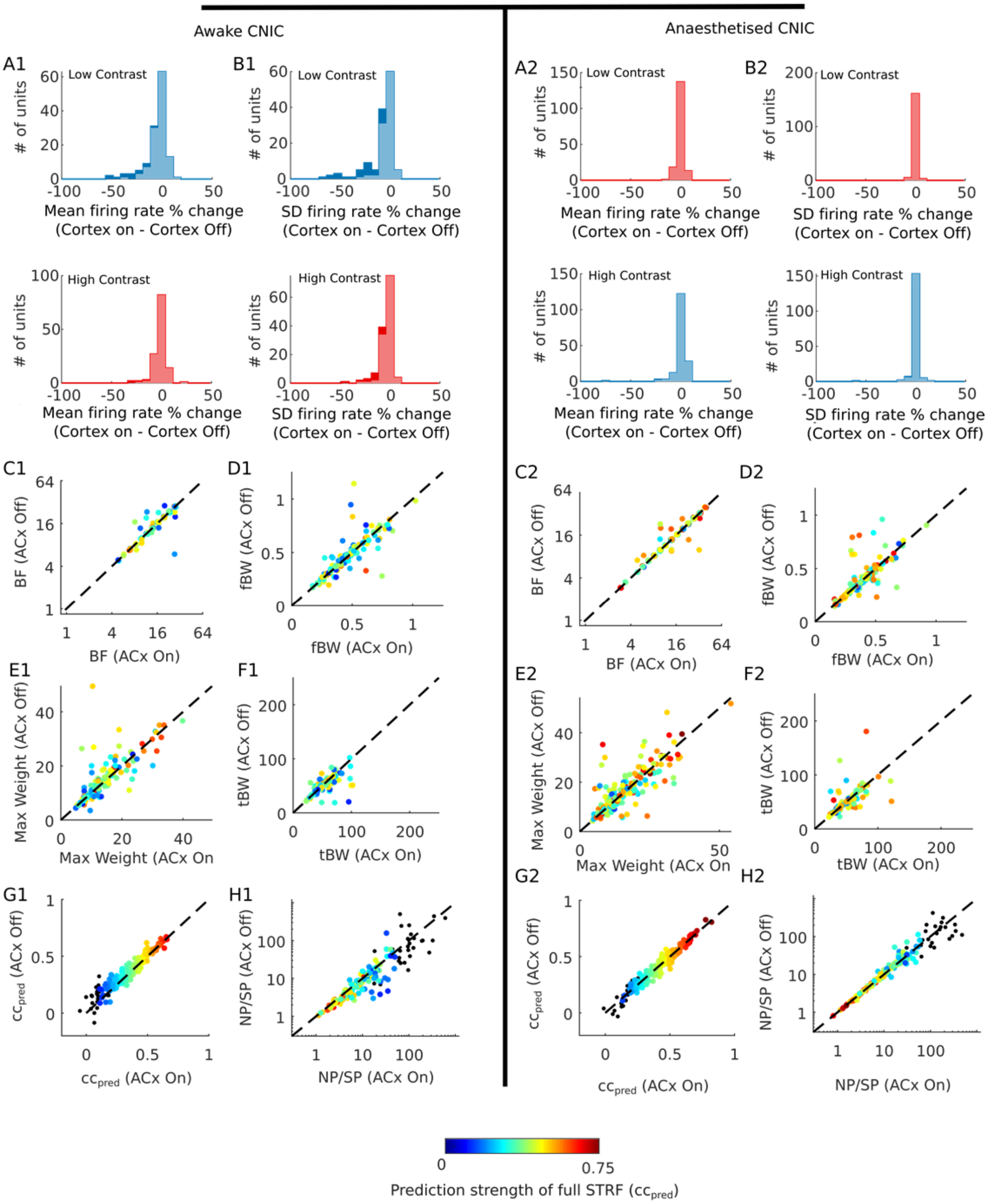


**Supplementary Figure 6. Cortical influences on responses to complex sounds in the inferior colliculus of awake and anesthetized mice.** (A) Change in mean firing rate in CNIC of awake (A1) and anesthetized (A2) mice during low contrast (top, blue) and high contrast (bottom, red) DRC stimulation following optogenetic cortical silencing. Wilcoxon signed rank tests: awake mice, $p_{Low\_Contrast} < 0.001$, $p_{High\_Contrast} = 0.003$; anesthetized mice, $p_{Low\_Contrast} > 0.5$, $p_{High\_Contrast} > 0.05$. (B) Change in standard deviation (SD) of firing rate across time in CNIC of awake (B1) and anesthetized (B2) mice during low contrast (top) and high contrast (bottom) DRC stimulation following optogenetic cortical silencing. Wilcoxon signed rank tests: awake mice, $p_{Low\_Contrast} < 0.001$, $p_{High\_Contrast} < 0.001$; anesthetized mice, $p_{Low\_Contrast} > 0.5$, $p_{High\_Contrast} > 0.05$. Light shaded areas in A and B indicate units that were not significantly modulated by cortical silencing, while dark areas represent units that were affected by cortical silencing ($p < 0.05$, t-test). (C) Comparison of the best frequency (BF), i.e., the largest value of the spectral kernel of the STRF, of CNIC units in awake (C1) and anesthetized (C2) mice between recordings made with auditory cortical activity intact (ACx On) or optogenetically silenced (ACx Off). (D) Frequency bandwidth (fBW), i.e., the full width half maximum (in octaves) around the BF, of CNIC units recorded in awake (D1) and anesthetized (D2) mice with and without cortical silencing. (E) Weight of the best frequency (BF weight) in the spectral kernel of the STRF of CNIC units recorded in awake (E1) and anesthetized (E2) mice with and without cortical silencing. (F) Temporal bandwidth (tBW), i.e., the full width half maximum (in ms) around the maximum weight of the temporal kernel of the STRF, of CNIC units recorded in awake (F1) and anesthetized (F2) mice with and without cortical silencing. (G) Linear model prediction performance within contrast (cross-validated correlation between predicted and actual responses) in the CNIC units of awake (G1) and anesthetized (G2) mice with and without cortical silencing. (H) NP/SP for CNIC units recorded in awake (G1) and anesthetized (G2) mice with and without cortical silencing (see Supplementary Methods). (C-H) Color of points denotes the prediction strength (correlation coefficient) of the model on a cross-validated dataset. Black dots are units excluded from analysis, according to exclusion criteria described in the STAR Methods. $n_{CNIC\_Anaesthetised} = 169$, 5 mice, $n_{\mathrm{CNIC\_Awake}} = 129$, 3 mice.

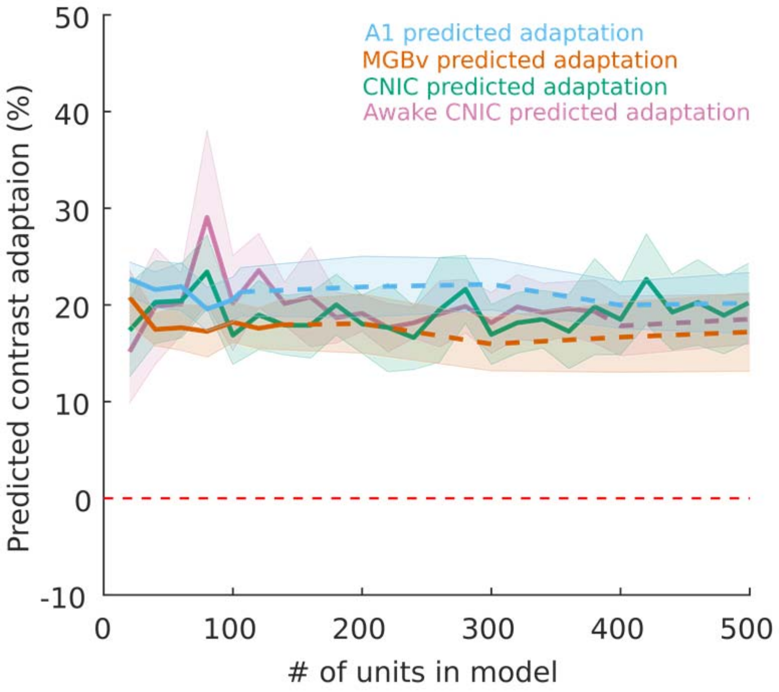


**Supplementary Figure 7. Predicted contrast adaptation as a function of number of units included in neurometric behavioral prediction model.** Solid lines indicate prediction of perceptual contrast adaptation where recorded units are used only once in each simulated trial. Dashed lines indicate prediction of perceptual contrast adaptation where recorded units are used more than once in each simulated trial (see Supplementary Methods). Shaded area denotes 95% confidence intervals of the means.

### Supplementary Methods

#### Measuring variance of neurons in the midbrain, thalamus, and cortex adapt to the contrast of the auditory environment – Analysis for Supplementary Figure 4

The contrast-specific LN models used in this paper to assess contrast adaptation require that units respond reliably to the DRCs and that those responses can be explained by a linear model. However, not all auditory neurons respond reliably and linearly to the DRCs. To probe contrast adaptation without having to exclude units based on their response reliability and linearity, we assessed whether neurons in the auditory pathway of mice can compensate for changes in contrast by adjusting their response variance. If the response of an auditory neuron matches the input, without adapting, it would be expected that its firing rate will display larger fluctuations from the baseline for high-contrast stimuli, which are characterized by larger level fluctuations, than for low-contrast stimuli. We can therefore determine whether the neuron displays contrast adaptation by measuring whether firing rate fluctuations decrease as expected when the contrast of the stimulus is reduced. Alternatively, if contrast adaptation is taking place, the response fluctuations will be more similar in high and low contrast conditions than expected.

To estimate the variance related to the input – and therefore whether the variance of the response adapts to the current auditory contrast – we employed a method proposed by Sahani and Linden (2003) to estimate variance related to the signal (sensory input) and the variance related to 'noise'. Following the terminology used by Sahani and Linden (2003), the variance of a response related to the input is called the *signal power*, and the variance of a response unrelated to the sensory input is called the *noise power*. The signal power is the variance related to the input because it is the variance of the mean firing rate over time relative to the variance seen in individual trials (i.e., total power).

$$SP = signal\ power = \frac{1}{N-1}\left(N \cdot \mathrm{Var}(y) - TP\right)$$

$$TP\ = total\ power = (N-1) \cdot \sum_{n=1}^{N} \mathrm{Var}(R_n)$$

To estimate whether the standard deviation related to the signal power adapts to contrast, we can take the ratio between $\sqrt{\text{signal power}}$ for low vs high contrast relative to the contrast change

$$signal\ ratio\ = \frac{s_{Low} C_{High}}{s_{High} C_{Low}}$$

$$s = \sqrt{signal\ power}$$

Where we double the contrast between low and high contrast, we can then assess the % contrast compensation in signal power using the same conversion as the gain ratio by

$$signal\ compensation = (signal\ ratio - 1) \times 100$$

#### Measuring dependence on unit numbers included in neurometric behavioral models – Analysis for Supplementary Figure 7

The behavioral prediction model presented in Figure 7 uses simulated responses from all individual units in each brain region and anesthetic state in each simulated trial. However, the number of units recorded in each case was different. In order to test the sensitivity of the model predictions to the number of units included in the model, we re-ran the model with different numbers of units included, by either down-sampling the number of units available (if using fewer units than available), or sampling with replacement if estimating the model with more units than were available from the recordings. The contrast adaptation prediction for each area was run 25 times (each consisting of 500 repeats per contrast condition).
